# A minimum reporting standard for multiple sequence alignments

**DOI:** 10.1101/2020.01.15.907733

**Authors:** Thomas KF Wong, Subha Kalyaanamoorthy, Karen Meusemann, David K Yeates, Bernhard Misof, Lars S Jermiin

## Abstract

Multiple sequence alignments (MSAs) play a pivotal role in studies of molecular sequence data, but nobody has developed a minimum reporting standard (MRS) to quantify the completeness of MSAs in terms of completely-specified nucleotides or amino acids. We present an MRS that relies on four simple completeness metrics. The metrics are implemented in AliStat, a program developed to support the MRS. A survey of published MSAs illustrates the benefits and unprecedented transparency offered by the MRS.

## INTRODUCTION

MSAs are widely used during annotation and comparison of molecular sequence data, allowing us to identify medically important substitutions (1), infer the evolution of species (2), detect lineage- and site-specific changes in the evolutionary processes (3) and engineer new enzymes (4). There is a wide range of computational tools for obtaining MSAs, and two of these (i.e., Clustal W (5) and Clustal X (6)) are now among the 100 most cited papers in science (7).

In addition to the completely specified nucleotides (i.e., A, C, G, T/U) or amino acids (i.e., A, C, D, E, F, G, H, I, K, L, M, N, P, Q, R, S, T, V, W, Y), MSAs may contain ambiguous characters (i.e., incompletely-specified nucleotides or amino acids). Frequently, they also contain alignment gaps (i.e., ‘–’) inserted between the nucleotides or amino acids of some of the sequences. Alignment gaps are inserted to maximize the homology of residues from different sequences (alignment gaps should only be used to improve alignment whereas N and X should only be used to signal missing data). A correct MSA is necessary for accurate genome annotation, phylogenetic inference and ancestral sequence reconstruction. However, deciding where to put the alignment gaps may be more art than science. This is because homology is defined as similarity due to historical relationships by descent (8). Most of these relationships belong to the unobservable distant past, so it is impossible to measure the accuracy of most MSAs inferred from real sequence data.

Without this ability, reporting the completeness of MSAs may be the best that can be achieved. So far, the only metric sometimes used is the *percent missing data* for a sequence (9) or an alignment (10), but neither is sufficiently transparent and informative. Recently, a guideline for systematic reporting of sequence alignments has been suggested (11), but it did not include completeness of MSAs—instead, it focused on quality indicators of alignment, but it did not define any of these or point to relevant literature. To rectify this, we developed a minimum reporting standard (MRS) for MSAs.

## MATERIALS AND METHODS

### Metrics for measuring completeness of MSAs

The MRS uses four metrics to quantify the completeness of different attributes of MSAs. Given an MSA with *m* sequences and *n* sites, we may compute four metrics: 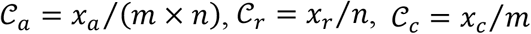 and 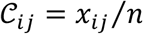, where *x_a_* is the number of completely specified characters (12) in the MSA, *x_r_* is the number of completely specified characters in the *r*th sequence of the MSA, *x_c_* is the number of completely specified characters in the *c*th column of the MSA and *x_ij_* is the number of homologous sites with completely specified characters in both sequences (*i* and *j*). In summary, 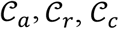 and 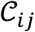 measure the completeness of the alignment, the *r*th sequence, the cth site, and the *i*th and *j*th sequences, respectively.

The first of these metrics 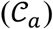 is related to the *percent missing data* used previously, but it is also, as shown in Figure 1A, the least useful completeness metric considered here: Alignments A and B differ greatly, but they have the same 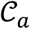 value (i.e., 0.7). The 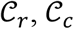 and 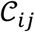 metrics, on the other hand, are able to detect these differences. For example, the 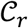 values range from 0.3 to 1.0 for Alignment A and from 0.4 to 1.0 for Alignment B, raising greater concern, from a sequence-centric perspective, about Alignment A than about Alignment B. If we were to omit any sequence from Alignment A, then it would be sensible to omit the one with the smallest 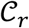 value. The 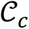 values range from 0.2 to 1.0 for Alignment A and from 0.5 to 0.8 for Alignment B. Again, there is greater concern about Alignment A than about Alignment B (due to the lower 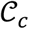 scores and the greater range of values). The 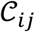 values range from 0.3 to 1.0 for Alignment A and from 0.0 to 0.9 for Alignment B. There is cause for great concern if 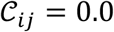 is detected as it means that sequences *i* and *j* have no shared homologous sites with completely specified characters in both sequences. Evolutionary distances between such sequences cannot be estimated unless the MSA contains at least one other sequence that overlaps both *i* and *j*. When such a case occurs, the evolutionary distance between sequences *i* and *j* is called *inferred by proxy*. Currently, the prevalence of this problem is unknown.

**Fig. 1.**
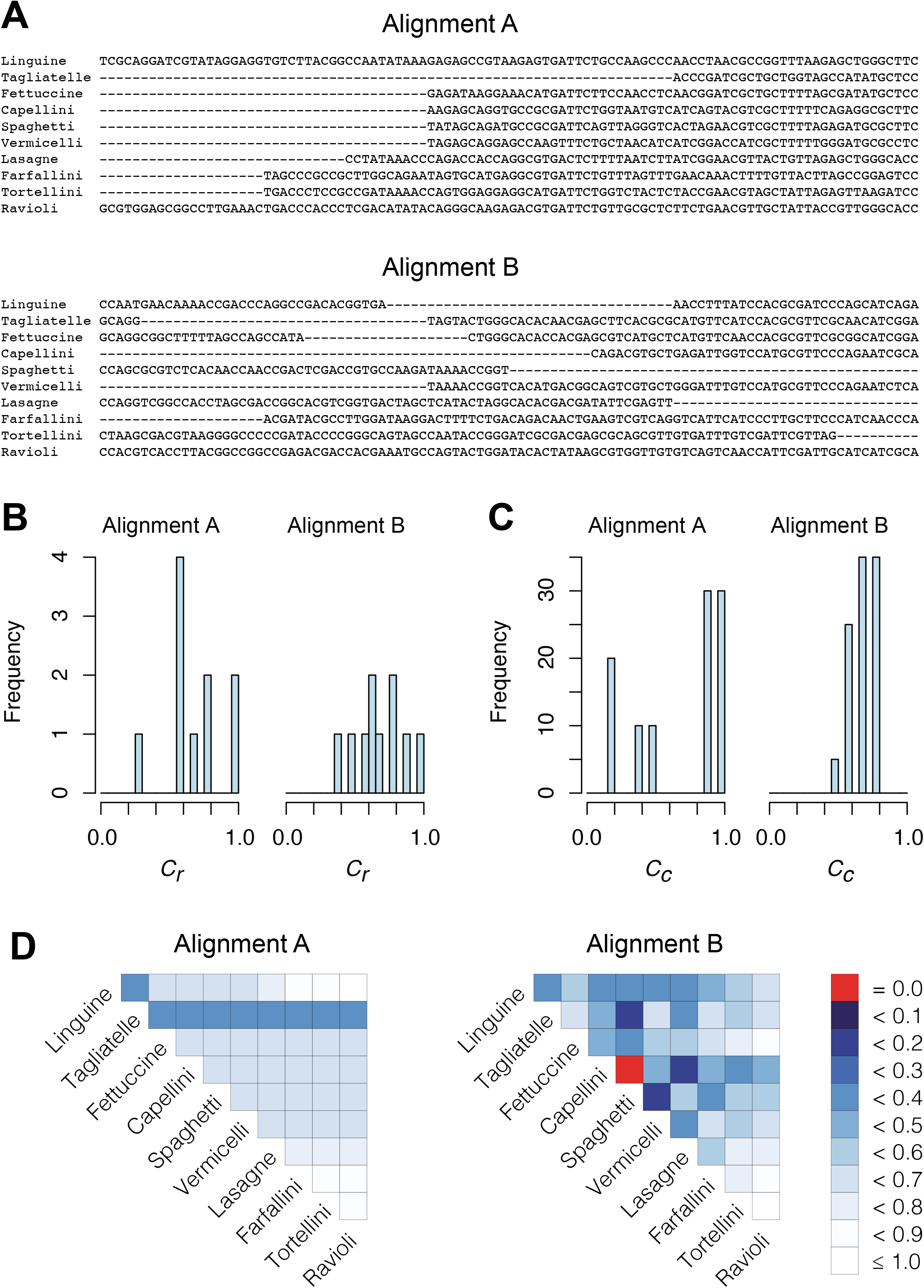
Example, based on two multiple sequences alignments (**A**), illustrating the corresponding distributions of completeness scores for rows (**B**), columns (**C**), and pairs of sequences (**D**).

Figures 1B and 1C reveal the distributions of 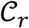 and 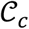 for Alignments A and B, offering additional insight into the alignments’ completeness. Conveniently, the 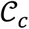 scores may be used to selectively omit the least complete sites. This *masking of sites* in MSAs is popular in phylogenetics and many methods (13–21) are now available. Additional information can be obtained by analyzing heat maps generated from the 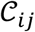 values. Figure 1D shows the heat maps obtained from Alignments A and B. The most obvious things to note are that in Alignment A *Tagliatelle* stands out as being the least complete sequence whereas *Capellini* and *Spaghetti* share no homologous sites with completely specified nucleotides in both sequences in Alignment B. Although this was easy to detect in Figure 1, it will be more difficult to do if *n* and/or *m* were larger, as is typically the case in phylogenomic data.

The benefits offered by the new completeness metrics are clear, but embedding figures like those in Figure 1 in publications may be impractical. Alternatively, the essential details may be reported in a table (table 1), or in one line (e.g., Alignment B: *m* = 10, *n* = 100, 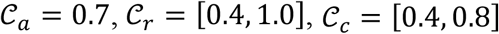, and 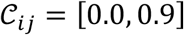). The closer to 1.0 the four 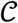 scores are, the more complete an alignment is. If, on the other hand, the values are closer to 0.0 than to 1.0, users may consider masking some of the sequences and/or sites before starting a phylogenetic analysis of the data.

**Table 1.** Example of the MRS for the alignments in Figure 1A

| Feature | Alignment A | Alignment B |
| --- | --- | --- |
| Sequences | 10 | 10 |
| Sites | 100 | 100 |
| Alphabet | Nucleotides | Nucleotides |
| $\mathcal{C}_a$ | 0.7 | 0.7 |
| $\mathcal{C}_r$ [min – max] | 0.3 – 1.0 | 0.4 – 1.0 |
| $\mathcal{C}_c$ [min – max] | 0.1 – 1.0 | 0.4 – 0.8 |
| $\mathcal{C}_{ij}$ [min – max] | 0.3 – 1.0 | 0.0 – 0.9 |

Given their potential to inform researchers across a wide range of scientific disciplines, we argue that *m, n*, 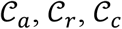 and 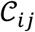 should be combined into what we henceforth call an MRS for MSAs, and that publications that report all of these values be labelled *compliant with the MRS for MSAs*. To our knowledge, this has never been done beforehand, leading to widespread ignorance about the MSAs that are relied upon in ground-breaking bio-medical research.

### AliStat: a program enabling compliance with the MRS for MSAs

To enable compliance with the MRS for MSAs, we developed AliStat, which is written in C++. To our knowledge, it is the first program to compute the four completeness scores presented above.

AliStat reads a text file with sequences of single nucleotides (i.e., a 4-state alphabet), di-nucleotides (i.e., a 16-state alphabet), codons (a 64-state alphabet) and amino acids (a 20-state alphabet), which are aligned and saved in the FASTA format. If the sequences comprise single nucleotides, then the characters may be ‘lumped’ to form six 3-state alphabets (i.e., CRT, AGY, ACK, GMT, AST and CGW) and seven 2-state alphabets (i.e., RY, KM, SW, AB, CD, GH and TV)—here R = A or G, Y = C or T, K = A or C, M = G or T, B = C or G or T, D = A or G or T, H = A or C or T, and V = A or C or G. If the 3- and 2-state alphabets are used, the letters R, Y, K, M, S, W, B, D, H and V are considered completely specified characters, unlike normal practice (12).

AliStat can be run in two modes: Brief mode or Full mode. Execution in brief mode is done using the following command:

~~~
alistat <infile> <data type> -b
~~~

and results in the following output format are printed to the terminal:

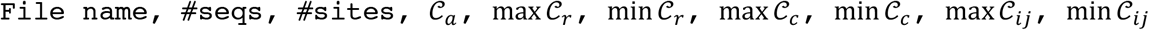

The brief-mode execution was included to allow users to quickly obtain the essential values from a great number of alignments (e.g., when comparing genomes phylogenetically).

The full-mode execution (default option) allows other options to be used and is intended when a more detailed examination of an MSA is required. For example, the -t option is used to indicate what types of 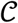 scores should be printed in output files, the -m option is used to set a threshold for masking sites, and the -i option is used to indicate that a heat map is needed. Other options and how all of the options may be used are described in the AliStat manual. The same information can be obtained by typing

~~~
alistat -h
~~~

in the command-line.

The output files appear in the .txt, .csv, .R, .dis, .svg, and .fst formats, which can be processed by other software packages. The .txt file summarizes the results. The .csv files present the 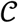 scores and may be examined using R. For example, if a user wishes to generate a histogram of the 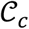 scores, the Table_2.csv file may be analyzed using the Histogram_Cr.R file. In some cases, users may want to infer a tree or network based on the 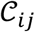 score (or the 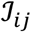 score, where 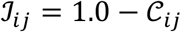). In such cases, .dis files may be analyzed by, for example, SplitsTree (22). The heat map, which may be triangular or square, is stored in the .svg file and may be opened using Adobe Illustrator™. If the - m option is used, the original MSA is split into two, with all sites having a 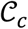 score larger than a user-specified threshold saved in a file called Mask.fst and the other sites saved in a file called Disc.fst. The two .fst files may be analyzed separately by other means (e.g., phylogenetic programs).

## RESULTS AND DISCUSSON

The MRS may be used to identify dubious MSAs. These alignments occur regularly in biomedical research and may also be present in large phylogenomic research, due to problems that might have arisen during the assembly, orthology assignment, and alignment procedures.

Typically, MSAs comprise more sequences and sites than those in Figure 1A, so to facilitate using the MRS, we implemented AliStat (Materials and Methods; http://github.com/thomaskf/AliStat), a fast, flexible, and user-friendly program for surveying MSAs. AliStat computes the 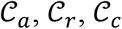, and 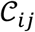 values from MSAs of nucleotides, di-nucleotides, codons and amino acids., AliStat lists the results on the command-line or in files that can be accessed by other programs.

The benefit of the MRS for MSAs is underlined in two surveys of large MSAs (table 2). In the first case, surveying an MSA of the enzyme carboxyl/cholinesterase (23) revealed that some of the 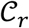 and 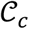 scores are closer to 0.0 than 1.0, and that at least two sequences have no homologous sites in common with completely specified characters in both sequences. Further inspection of the output files revealed large proportions of low 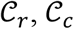, and 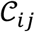 scores (Figs, S1, S2, and S3; Supplementary Material), so it might be wise to mask some of the sequences or sites before phylogenetic analysis of these data. Given the main objective of the original analysis of these data (to annotate the genes in two major crop pests), masking sites with completeness scores below 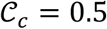 had a big impact on the 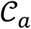 score (it increased from 0.2262 to 0.9562) and, hence, also on the maximum scores of 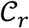 and 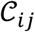 (Table S1; Supplementary material).

**Table 2.** Example of the MRS for two published MSAs

| Feature | Carboxyl/Colineesterase<br>(23) | Lepidoptera (24) |
| --- | --- | --- |
| Sequences | 364 | 203 |
| Sites | 2,645 | 749,791 |
| Alphabet | Amino acids | Amino acids |
| $\mathcal{C}_a$ | 0.2262 | 0.6422 |
| $\mathcal{C}_r$ [min – max] | 0.0106 – 0.5550 | 0.0609 – 0.9738 |
| $\mathcal{C}_c$ [min – max] | 0.0027 – 0.9972 | 0.0000 – 0.9655 |
| $\mathcal{C}_{ij}$ [min – max] | 0.0000 – 0.5550 | 0.0084 – 0.9672 |

In the second case, surveying a massive concatenation of MSAs of nuclear genes (24) revealed a more complete alignment but also low 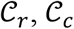 and 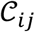 values. The presence of these values shows that additional masking of this MSA might have been wise (Figs. S4, S5, and S6; Supplementary Material). For example, omitting the two most incomplete sequences (i.e., the genera *Leucoptera* and *Pseudopostega*) could have been considered (Figs. S4 and S6; Supplementary Material).

The MRS for MSAs is a robust solution to a large and so-far-neglected problem: how do we report, as transparently and informatively as possible, the completeness of the MSAs used in bio-medical research? Better transparency about the completeness of MSAs is clearly needed, because MSAs represent a foundational cornerstone in many bio-medical research projects and, as revealed by the example in Figure 1, MSAs may look different but have the same percentage of missing data. So far, information on the completeness of MSAs used in bio-medical research has been largely absent, leaving readers unable to critically evaluate the merits of scientific discoveries made on the basis of MSAs. It is critical to recognize, and acknowledge, that many MSAs are the result of scientific procedures. Therefore, it is necessary to present the results of these procedures more transparently and comprehensively. Many scientific papers now include links to the MSAs used, but the MSAs are often so large that it is impossible to form a comprehensive picture about the completeness of these MSAs.

Our MRS enables a radical change in scientific behavior, allowing authors to report their results more transparently, and readers the ability to critically assess discoveries made from analyses of sequence data stored in MSAs.

## AVAILABILITY

AliStat is available from http://github.com/thomaskf/AliStat/ under an CSIRO Open Source Software License Agreement (variation of the BSD / MIT License).

## SUPPLEMENTARY DATA

Supplementary Material is available from NARGAB Online (see attachment).

## ACKNOWLEDGEMENTS

We thank staff at the Australian National University and University College Dublin for feedback on the color scheme used in the heat map; many of the respondents are color-blind. Finally, we wish to thank three reviewers for their constructive comments.

## Supplementary Material

### Analysis of an alignment of carboxyl/cholinesterases (CCEs) from a paper by Pearce et al. (1)

The alignment of amino acids used to annotate the CCE genes from *Helicoverpa armigera, H. zea, Manduca sexta* and *Bombyx mori* was surveyed using AliStat v1.11. The alignment comprised 364 sequences and 2645 sites. Figure S1 presents the distribution of 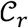 values, with several sequences having values close to 0.0. Because the objective of the study by Pearce et al. (1) was to annotate the genes, it was not possible to remove any sequences from the data.

**Figure S1.**
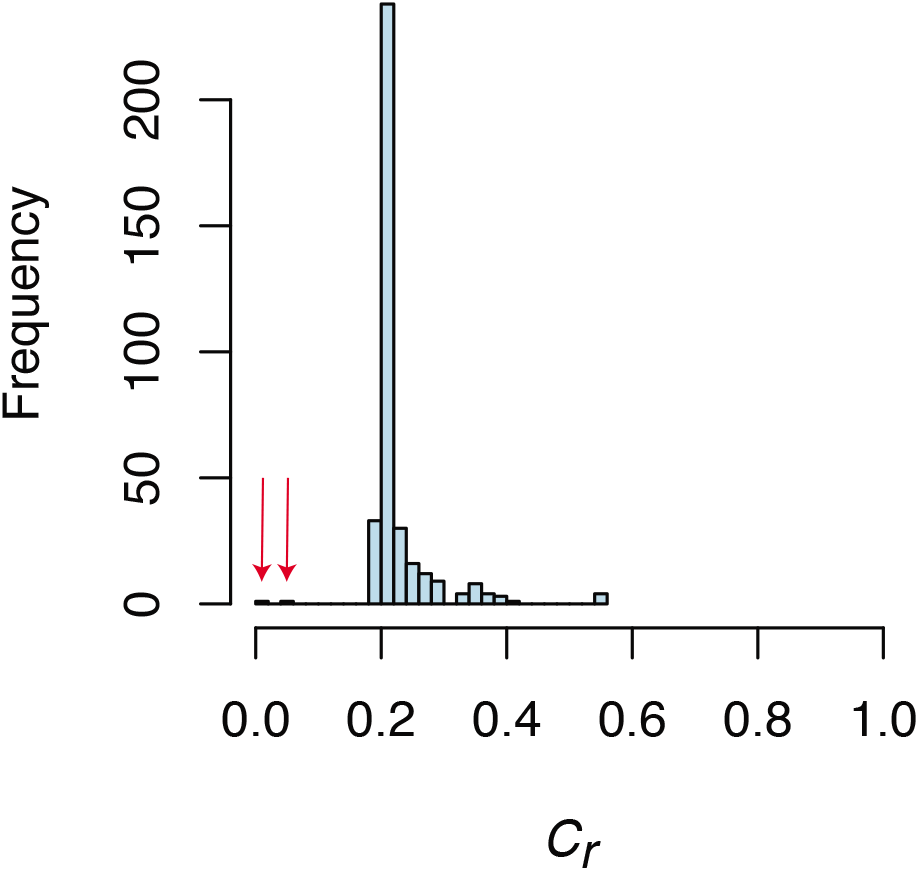
Histogram showing the distribution of 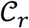 scores from the CCE genes. The arrows point to the lowest 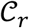 scores.

Figure S2 reveals the distribution of 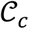 scores, with a high proportion of sites with low 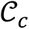 values.

Based on this distribution, sites with 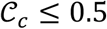 were masked in the study by Pearce et al. (1).

**Figure S2.**
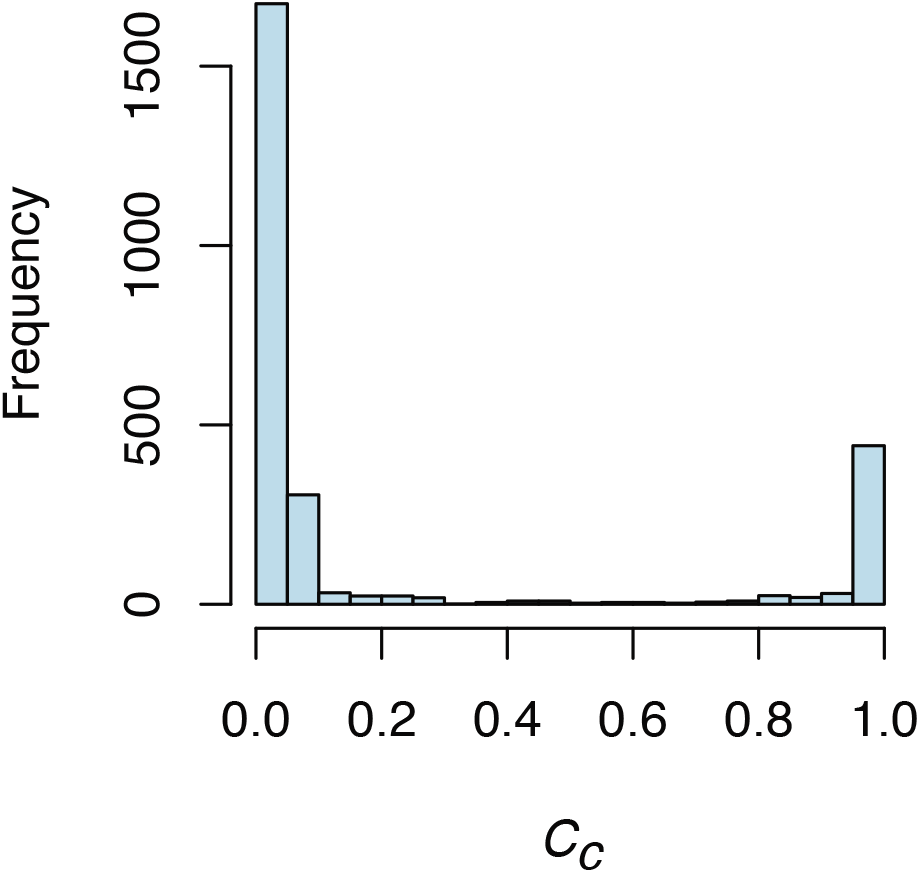
Histogram showing the distribution of 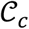 scores from the CCE genes.

Figure S3 shows the heat map of 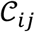 scores, revealing a high proportion of sequence pairs with low 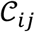 values. Because the aim of the study by Pearce et al. (1) was to annotate the genes, it was not possible to remove any sequences from the data.

**Figure S3.**
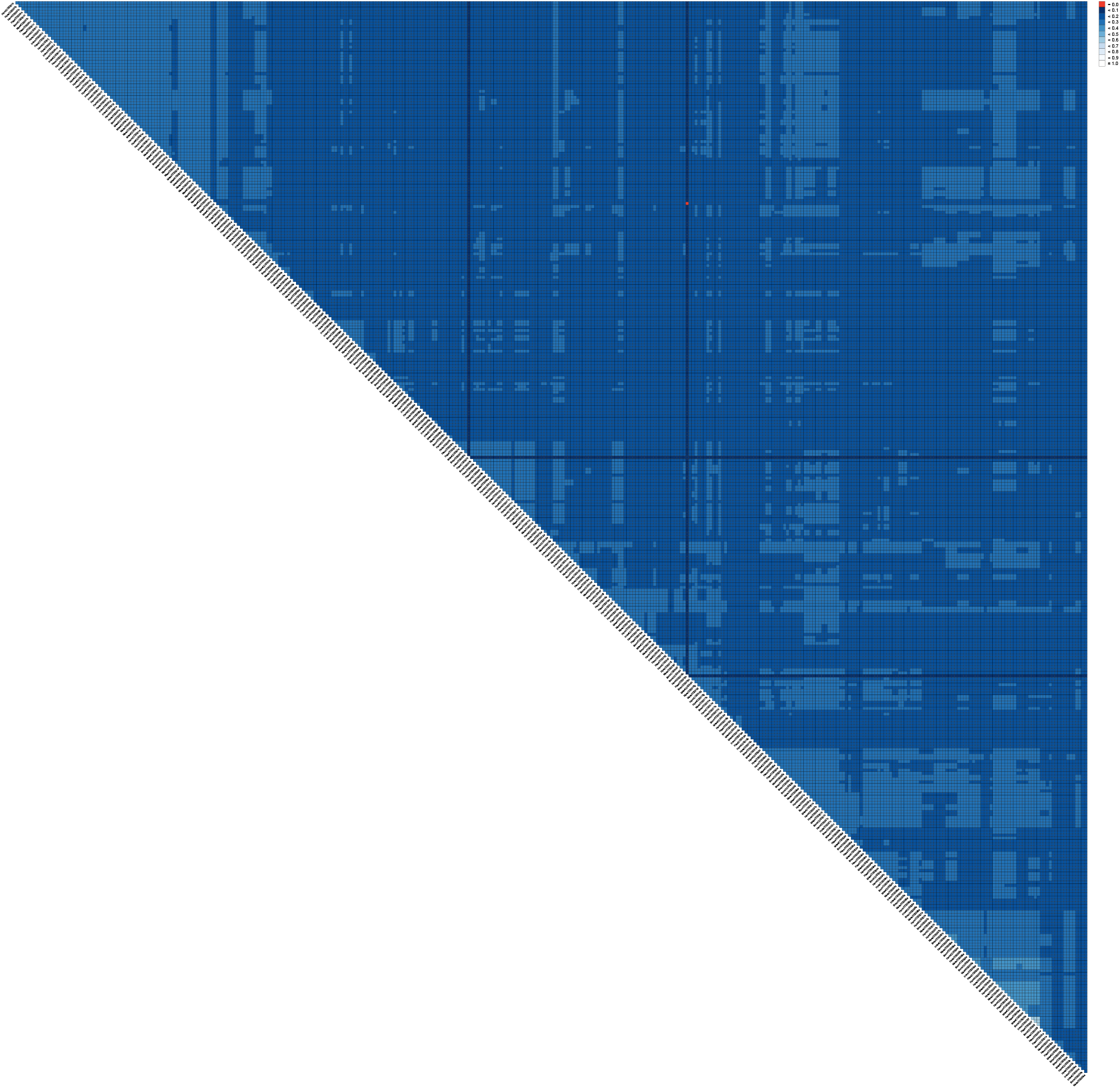
Heat map showing the distribution of 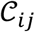 scores from the CCE genes. The benefit of this heat map is realized by enlarging the image, at which point it become obvious that two sequences, labelled HzeaCCE016h and MsexCCE001s, have no homologous sites with unambiguous characters in both sequences.

Pearce et al. (1) used a threshold of 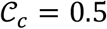 to mask the alignment of amino acids. Table 1 reveals the effect of doing so. In this case, the table complies with the minimum reporting standard (MRS) for multiple sequence alignments (MSAs).

**Table S1.** Example highlighting the effect of using 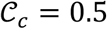 to mask the sites in the alignment of CCEs.

| Feature | Before masking | After masking |
| --- | --- | --- |
| Sequences | 364 | 364 |
| Sites | 2,645 | 546 |
| Alphabet | Amino acids | Amino acids |
| $C_a$ | 0.2262 | 0.9562 |
| $C_r$ [min – max] | 0.0106 – 0.5550 | 0.0458 – 0.9890 |
| $C_c$ [min – max] | 0.0027 – 0.9972 | 0.5055 – 0.9972 |
| $C_{ij}$ [min – max] | 0.0000 – 0.5550 | 0.0000 – 0.9890 |

### Analysis of lepidopteran nuclear data from a study by Kawahara et al. (2)

The alignment of amino acids labelled SA4_aminoacid_supermatrix_resorted_renamed.fas was surveyed using AliStat v1.11. The alignment comprised 203 sequences and 749,791 sites. Figure S4 reveals the distribution of 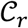 values, with most sequences having values close to 1.0. In this case, only a few sequences had very low 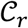 scores.

**Figure S4.**
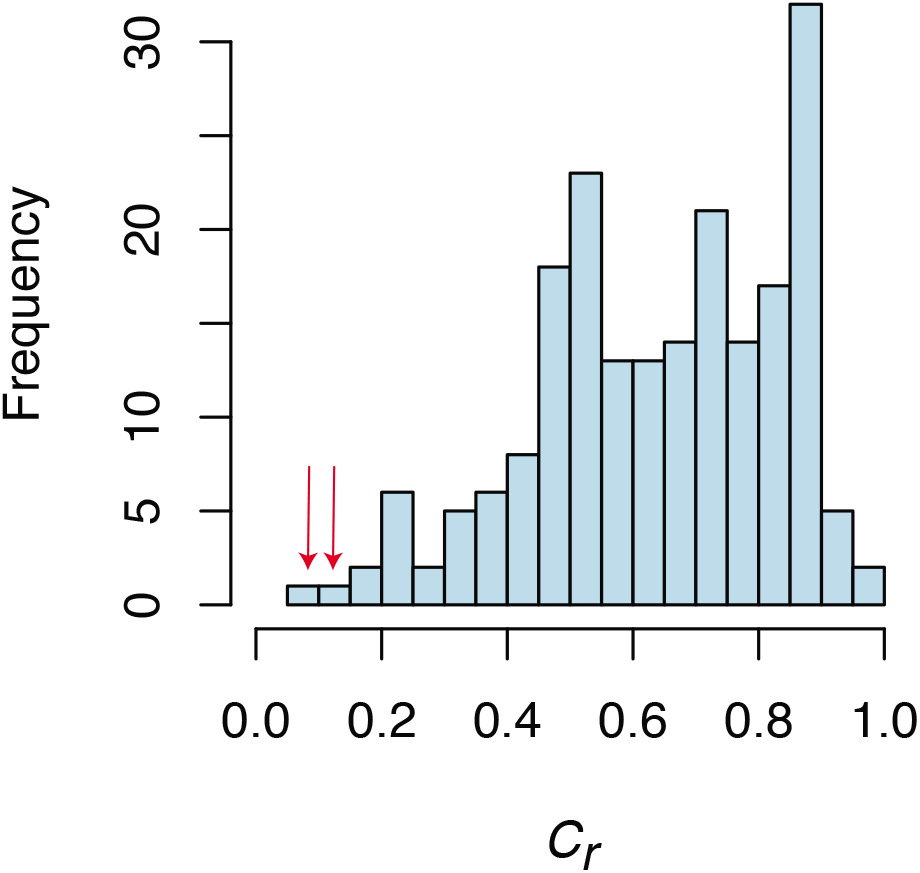
Histogram showing the distribution of 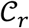 scores from the super-alignment (i.e., 2098 MSAs of single-copy genes concatenated). The arrows point to the two lowest 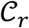 scores (genera *Leucoptera* and *Pseudopostega*).

Figure S5 reveals the distribution of 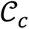 scores, with a high proportion of sites with high 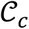 values. Based on this distribution, omitting sites with 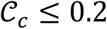 might have been sufficient.

**Figure S5.**
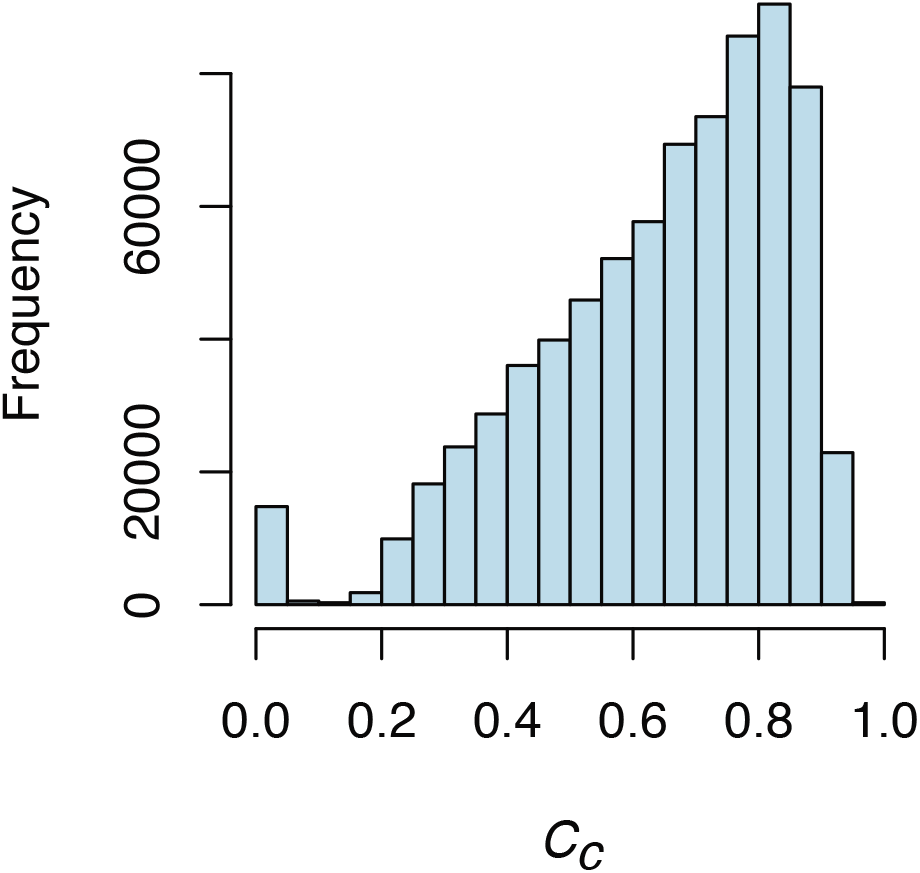
Histogram showing the distribution of 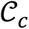 scores from the super-alignment (Kawahara et al. (2)).

Figure S6 shows the heat map of 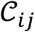 scores, revealing a low proportion of sequence pairs with low 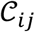 values. In this case, the sequences found to be least complete are from the genera *Leucoptera* and *Pseudopostega*.

**Figure S6.**
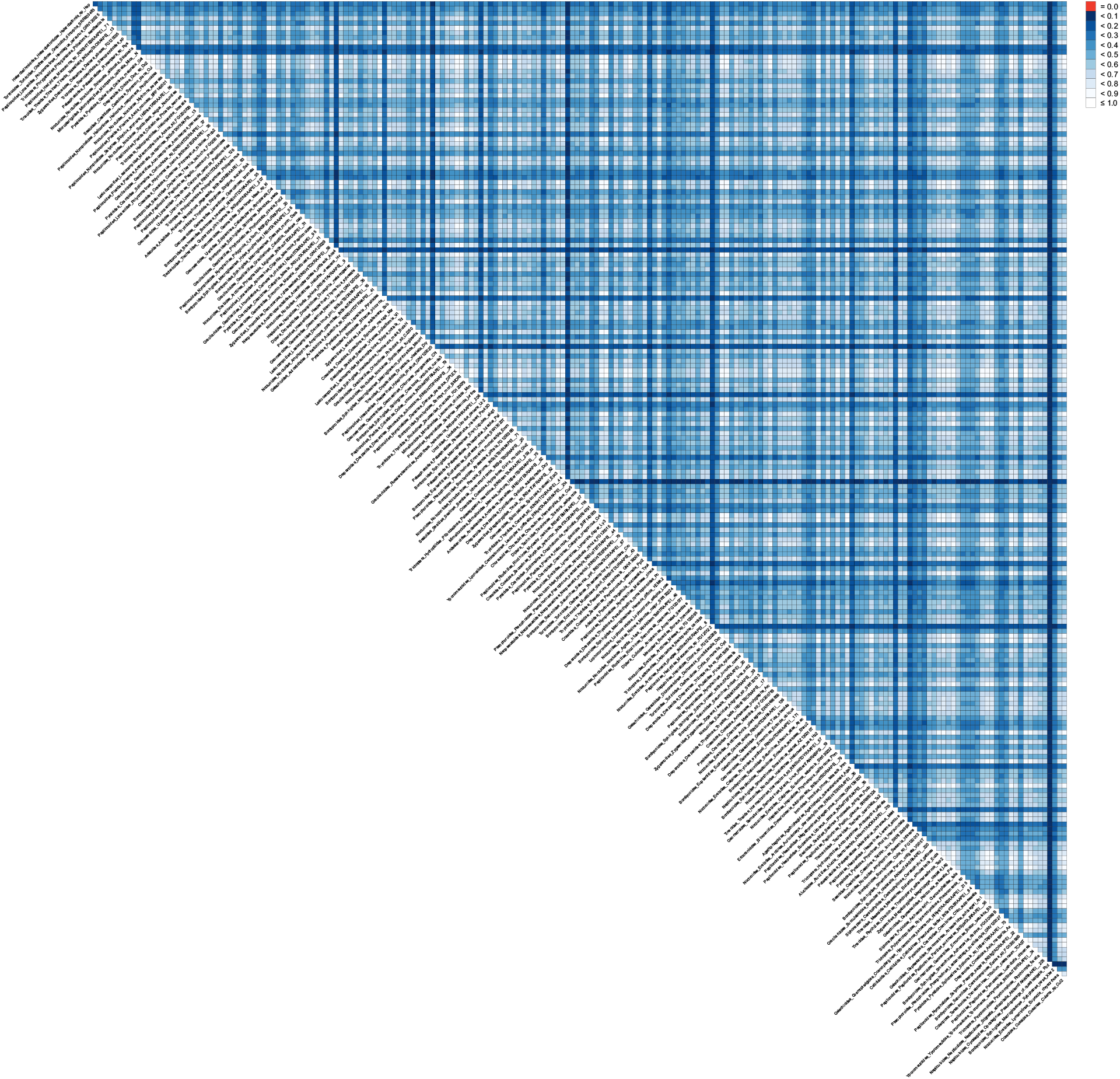
Heat map showing the distribution of ĉ¿$scores from the concatenate gene alignments. Again, the benefit of this heat map is realized by enlarging the image. In this case, the two most incomplete sequences are identified as being from the genera *Leucoptera* and *Pseudopostega*.

